# Latitude influences stability via stabilizing mechanisms in naturally-assembled forest ecosystems at different spatial grains

**DOI:** 10.1101/2022.12.07.519439

**Authors:** Tianyang Zhou, Dylan Craven, Holger Kreft, Jiaxin Zhang, Xugao Wang, Juyu Lian, Wanhui Ye, Guangze Jin, Xiangcheng Mi, Yujun Yao, Haibao Ren, Jie Yang, Min Cao, Congrong Wang, Yuanzhi Qin, Gang Zhou, Yanan Fei, Yaozhan Xu, Xiujuan Qiao, Mingxi Jiang, Nathaly R. Guerrero-Ramírez

**Author notes:** Corresponding author: Xiujuan Qiao.

## Abstract

Ecosystem stability reveals how ecosystems respond to global change over time. Yet, the focus of past research on small spatial grains and extents overlooks scale dependence and how broad-scale environmental gradients shape stability. Here, we use forest inventory data covering a broad latitudinal gradient from the temperate to the tropical zone to examine cross-scale variation in stability of aboveground biomass and underlying stabilizing mechanisms. While stability did not shift systematically with latitude at either spatial grain, we found evidence that species asynchrony increased towards the tropics at the small spatial grain while species stability decreased at both spatial grains. Moreover, latitude stabilized forest communities *via* its effects on both stabilizing mechanisms, which compensated for the weak and destabilizing effects of species richness. Yet, the trade-off in the relative importance of species stability and species asynchrony for stability was not mediated by latitude, suggesting that context-dependent factors - to a greater extent than macroecological ones - underlie large-scale patterns of stability. Our results highlight the crucial role of species asynchrony and species stability in determining ecosystem stability across broad-scale environmental gradients, suggesting that conserving biodiversity alone may not be sufficient for stabilizing naturally-assembled forest ecosystems.

## Introduction

Given the evidence that diverse communities can buffer environmental fluctuation to a greater extent than less diverse communities^1–4^ and thus stabilize ecosystem function over time (i.e., temporal stability or invariability), the diversity-stability relationship has been one of core research issues in ecology. The stabilizing effects of biodiversity on ecosystem functioning emerge largely due to two complementary, and not mutually exclusive, ecological mechanisms: species asynchrony and species stability^5^. Species asynchrony captures the heterogeneity in the response of co-occurring species to similar environmental conditions^6,7^, whereas species stability reflects the stabilizing effects of dominant species on temporal stability, whose impacts may – or may not – be mediated by biodiversity^8–10^. Yet, our understanding of diversity-stability relationships in naturally-assembled forest ecosystems comes mostly from studies with a limited geographic extent (i.e., the total area covered by all sampling units; but see: ^11,12^) and small spatial grains (i.e., the spatial scale of sampling; average spatial grain (or plot size) = 0.04 ha^13–15^). This limits our understanding of how temporal stability and its drivers vary across both aspects of spatial scale, and it remains uncertain whether results from small spatial grains and extents can provide a robust theoretical basis for developing management policies that target large areas for forest conservation.

With increasing spatial grain, the range and heterogeneity in environmental conditions are expected to increase^16^, resulting in a strengthening of biodiversity-ecosystem functioning (BEF) and diversity-stability relationships. For instance, stronger positive biodiversity effects on ecosystem functioning are expected at larger spatial grains since higher environmental heterogeneity and a broader range of environmental conditions provide more opportunities for niche partitioning^17^. Yet, an analysis of 25 naturally-assembled forests^18^ found that positive BEF relationships emerged mainly at the smallest spatial grain (0.04 ha), while BEF relationships shifted in direction to either neutral or negative at the largest spatial grain (1 ha). However, recent studies reported that biodiversity at large spatial grains could also enhance and stabilize ecosystem functions due to ‘macroecological complementarity’ effects, i.e. that diverse communities’ ability to buffer against environmental fluctuations should be more pronounced at larger spatial grains because more species with different ecological niches accumulate, or ‘spatial insurance’ effects, i.e. higher spatial turnover of biodiversity can stabilize regional community dynamics, implying that similar ecological processes may occur at multiple spatial scales^19–23^. As such, whether the current understanding of stability from local spatial grains can predict stability patterns and underlying stabilizing mechanisms at larger spatial grains – and whether they generate emergent macroecological patterns – is unclear.

Macroecological patterns of stability and shifts in stabilizing mechanisms may emerge due to changes in species richness, tree community properties (i.e. functional composition and stand structure), and environmental conditions. For instance, due to the stabilizing effects of biodiversity, it is expected that stability will change with latitude as is typically the case for species richness, i.e., increasing from the poles towards the equator^24^. Additionally, because forests can be less even towards the equator^25^, ecosystem functioning at higher latitudes may be regulated more strongly by dominant species^26,27^. Consequently, the stabilizing effects of species stability may be stronger at higher latitudes, while species asynchrony may follow the inverse pattern. Beyond biotic factors, latitudinal variation in abiotic factors, such as climate, have been shown to explain not only ecosystem stability^28,29^, but also the diversity-stability relationship^30,31^. Variation in climate regulates the spatial pattern of functional composition^32^, thus influencing ecosystem functions (as predicted by the mass ratio hypothesis^33^), such that communities with high biomass stocks and stability are typically dominated by species with trait values associated with resource conservation^33–36^. Similarly, stem density also varies with climate^12,13^, and often reflects interactions among individuals within a community, which in turn could influence temporal stability. In areas with fertile soil and warm climate, competition could destabilize ecosystem functioning, while in areas with barren soil and cold climate facilitation could increase temporal stability^13,37^.

To elucidate large-scale patterns of temporal stability and underlying stabilizing mechanisms in naturally-assembled forests, we analyzed a comprehensive census of six plots covering a broad latitudinal gradient (21.61 °N - 47.18 °N) ranging from temperate to tropical forests in the Chinese Forest Biodiversity Monitoring Network over ten years (Fig. 1A). Our dataset contains more than 1,900,000 re-measurements over three censuses with 1,013 species from these six plots, whose area range from 9 ha to 25 ha and are divided in quadrants of 0.4 ha (Table S1). To account for differences in plot size, we randomly selected 100 quadrant per plot (4.0 ha) 100 times (Fig S1). We address the following questions: (1) How do stability and stabilizing mechanisms (i.e. species asynchrony and species stability) vary with latitude, and how do latitudinal patterns vary with spatial grain (0.4 ha and 4.0 ha)? (2) What are the principal drivers that underpin stability? (3) Is there a latitudinal trade-off between species stability and species asynchrony in driving ecosystem stability? We hypothesize (H1) that the latitudinal pattern of stability may be either positive, negative, or neutral (Fig. 1B) due to three different opposing mechanisms. A positive relationship may emerge given the potentially destabilizing effects of warmer climates, resulting in stability increasing with latitude^28,29,38^. A negative relationship may emerge if the stabilizing role of biodiversity is stronger than the destabilizing effects of climate, mirroring the latitudinal diversity gradient^39^, and resulting in stability decreasing with latitude; or there might not be a directional change in stability with latitude, as biodiversity and stability may be decoupled in naturally-assembled ecosystems^36,40^. We anticipate that species asynchrony and species stability will decrease and increase with latitude, respectively, and that their stabilizing effects will be stronger at the smaller spatial grain, with weak or neutral effects at the large grain (Fig. 1C). Second, we hypothesize (H2) that both biotic (species richness, species stability, species asynchrony, functional trait composition, and stem density) and abiotic factors (latitude, climate and soil nutrients), directly and indirectly, influence variation in temporal stability across the latitudinal gradient. Finally, while we expect (H3) that both species asynchrony and stability will jointly determine stability, we anticipate that their relative importance will exhibit a trade-off along the latitudinal gradient (Fig. 1D), with stability mainly being driven by species stability at higher latitudes and by species asynchrony at lower latitudes.

**Fig. 1.**
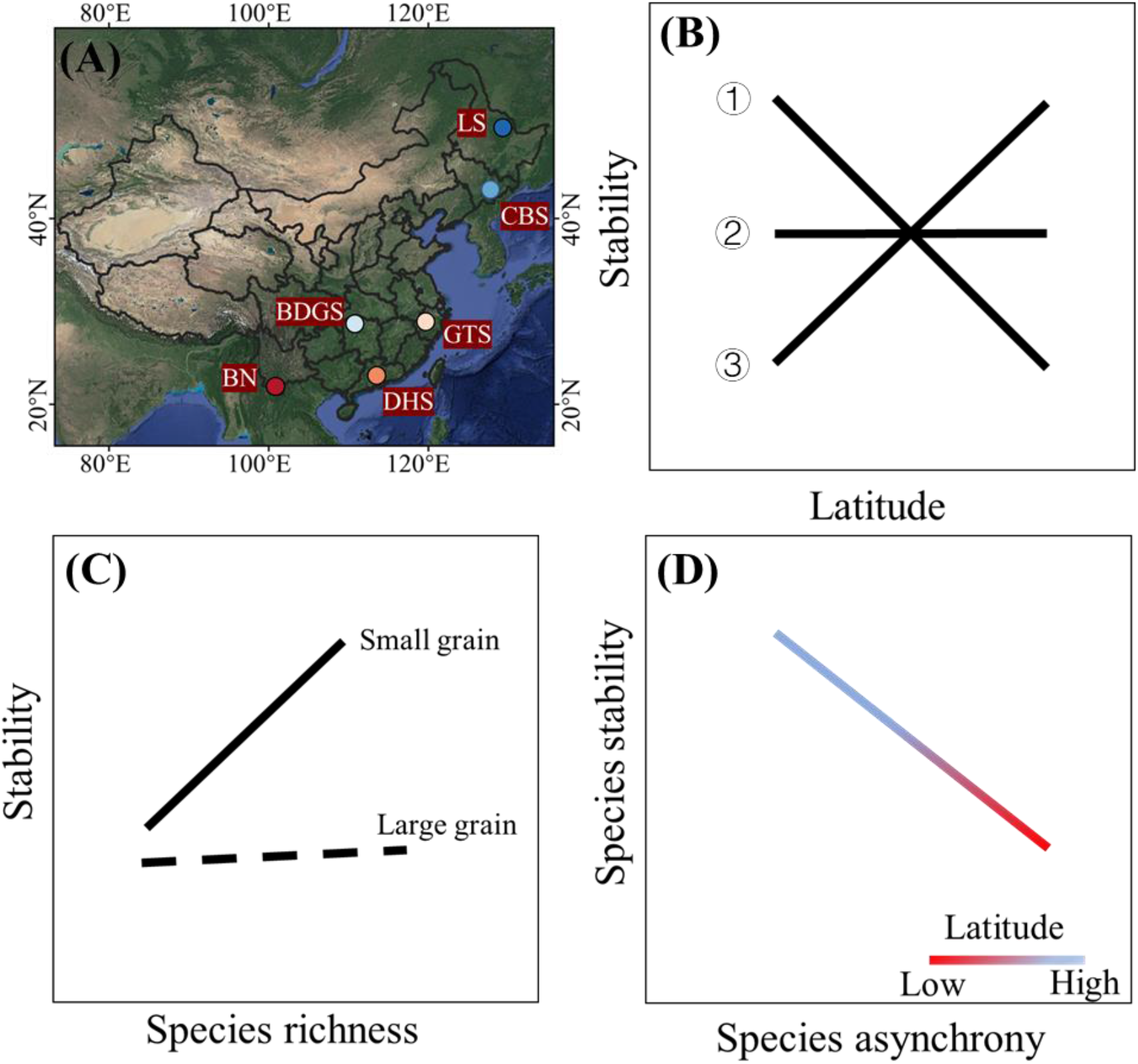
The latitudinal gradient used in this study to examine broad-scale patterns and the hypothesized patterns, drivers of ecosystem stability. (A) The spatial distribution of six study sites belonging to three forest ecosystems across China (temperate forest: LS (Liangshui) and CBS (Changbaishan); subtropical forest: BDGS (Badagongshan), GTS (Gutianshan), and DHS (Dinghushan); tropical forest: BN (Banna), Table S1). (B) Hypothesized latitudinal patterns of stability. Given a stabilizing effect of biodiversity, we would expect that stability is higher at lower latitudes, and decreases with increasing latitude (1). However, biodiversity and stability could be decoupled in naturally-assembled ecosystems, resulting in a non-directional relationship between stability and latitude (2). Due to the destabilizing effects of warmer climates, stability also could decrease with latitudinal shifts in temperature (3). (C) Hypothesized relationships between species richness and stability at different spatial grains. The positive effects of species richness on stability at small spatial grain (0.04 ha) could weaken with spatial grain, possibly becoming neutral at the large spatial grain (4.0 ha). (D) The prediction that the relative importance of species stability and asynchrony for ecosystem stability in forest communities may vary across the latitudinal gradient. Species asynchrony could play a more important role in shaping the stability of tropical forests due to their higher biodiversity, while species stability could play a more significant role in determining the stability of temperate forests, which are frequently dominated by fewer species (i.e. they are less even) than tropical forests.

## Results

### Latitudinal patterns in temporal stability and stabilizing mechanisms across spatial grains

Our results indicated that stability did not vary significantly with latitude at either spatial grain (large grain: *P* = 0.08, small grain: *P* = 0.13, Fig. 2A, D). In contrast, we found that species asynchrony significantly decreased with latitude at the small, but not at the large spatial grain (large grain: *P* = 0.61; small grain: *R^2^*m = 0.12, *P* = 0.04, Fig. 2B, E) and that species stability significantly increased with latitude at both spatial grains (large grain: *R^2^*m = 0.78, *P* < 0.01; small grain: *R^2^*m = 0.27, *P* < 0.01, Fig. 2C, F).

**Fig. 2.**
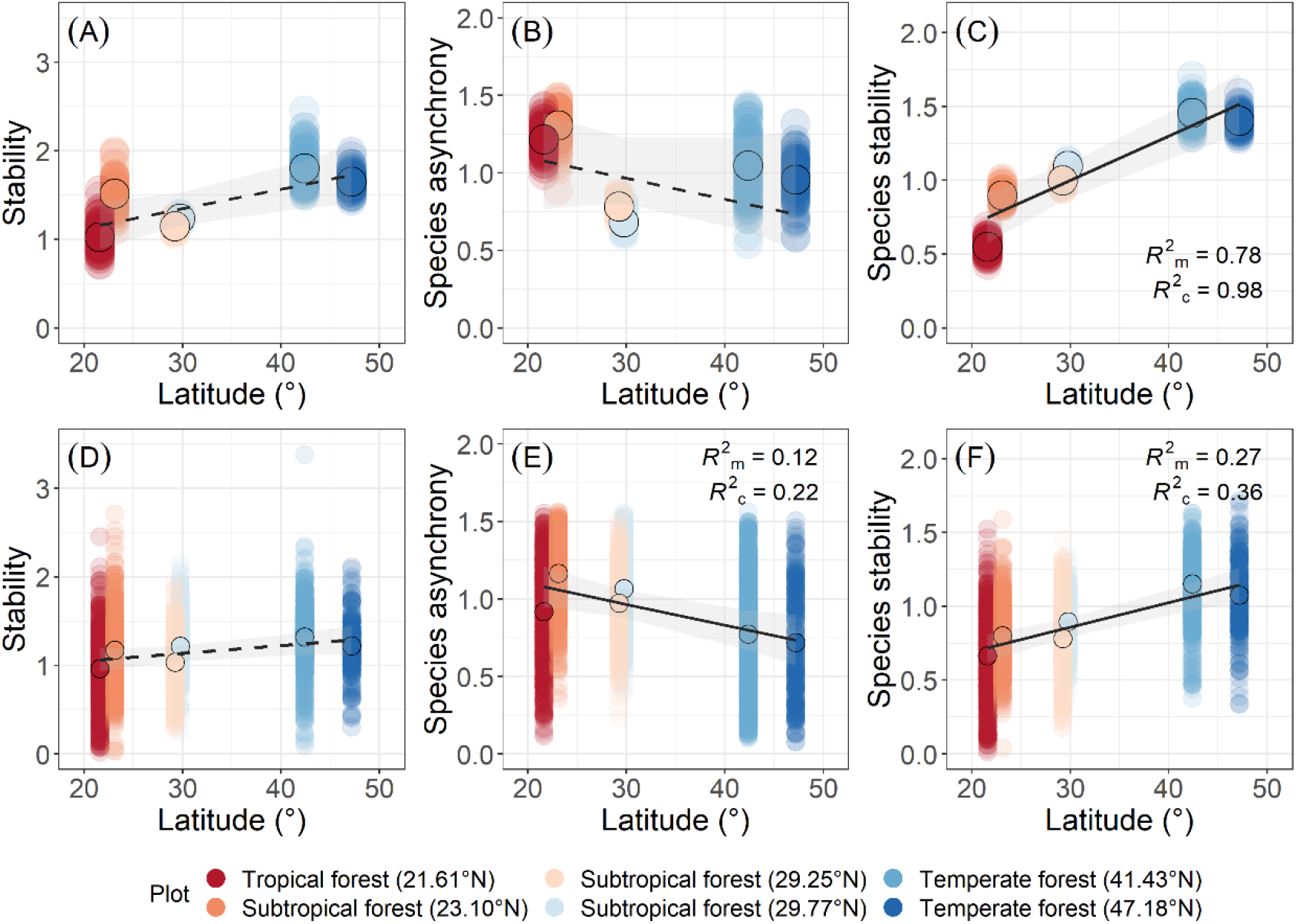
Latitudinal patterns of stability (A, D), species asynchrony (B, E) and species stability (C, F) at two spatial grains (4.0 ha (n = 600) and 0.4 ha (n = 3075)). Lines are mixed-effects model fits (solid lines: *P* < 0.05; dashed line: *P* > 0.05). Translucent points are plot-level values, while opaque points with black circles are mean values for each of the six study sites. Different colors indicate different study sites. *R^2^*m and *R^2^*c represent variation explained by fixed effects and the combination of fixed and random effects in mixed-effects models, respectively. Grey bands represent 95% confidence intervals. Both stability and species stability were log10 transformed, and species asynchrony was angular transformed.

Our results showed that species richness destabilized forest communities at both spatial grains (large grain: *R^2^*m = 0.53; small grain: *R^2^*m = 0.03; Fig. 3A, D; Table S7), while species asynchrony (large grain: *R^2^*m = 0.30; small grain: *R^2^*m = 0.32; Fig. 3B, E) and species stability stabilized forest communities at both spatial grains (large grain: *R^2^*m = 0.67; small grain: *R^2^*m = 0.45; Fig. 3C, F; Table S7).

**Fig. 3.**
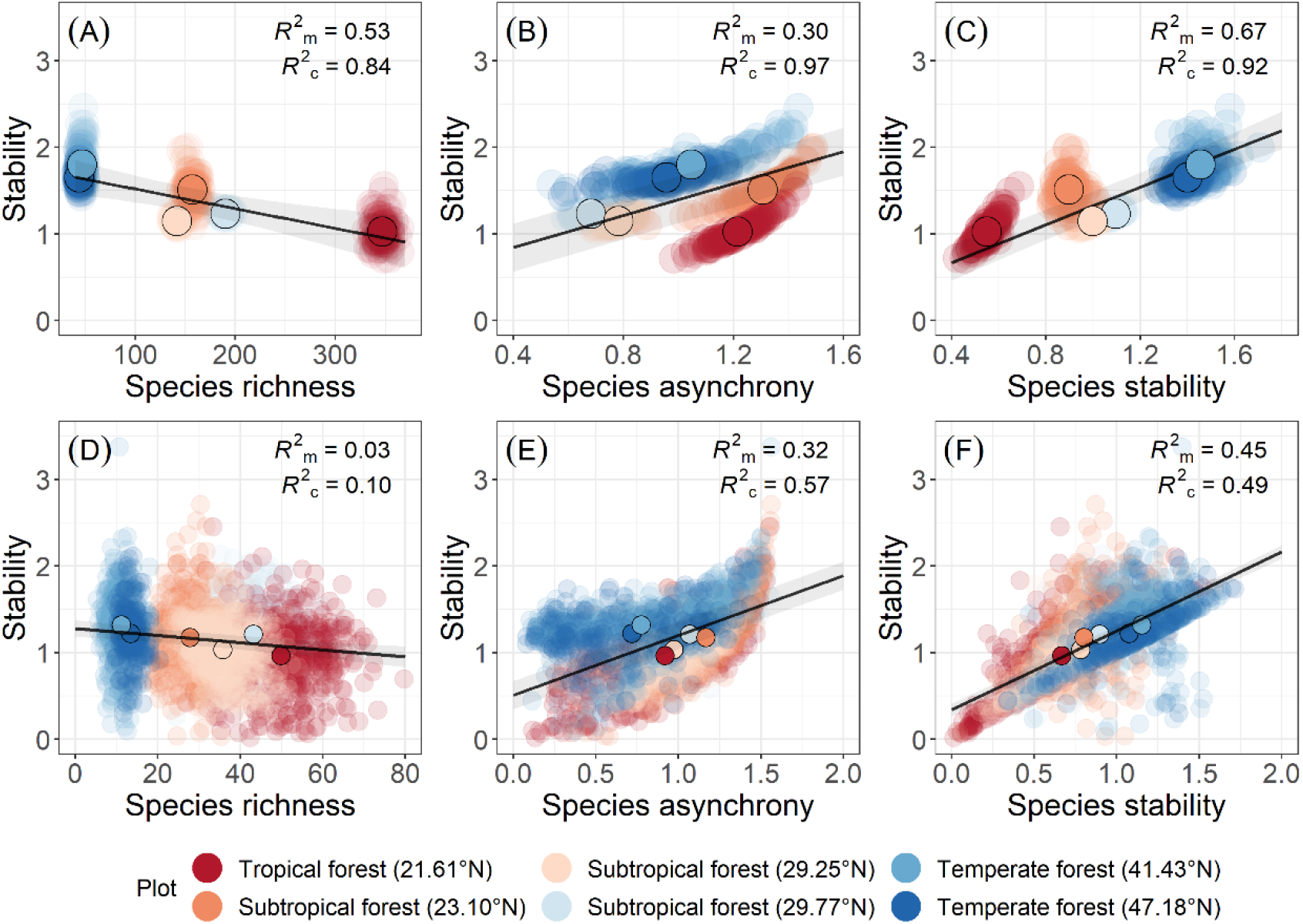
Relationships between species richness (A, D), species asynchrony (B, E), species stability (C, F) and stability at two spatial grains (4.0 ha (n = 600) and 0.4 ha (n = 3075)). Lines are mixed-effects model fits. Translucent points are plot-level values, while opaque points with black circles are mean values for each study site. Different colors indicate different study sites. *R^2^_m_* and *R^2^_c_* refer to the marginal and conditional *R^2^*, which represent variation explained by fixed effects and the combination of fixed and random effects in mixed-effects models, respectively. Light grey bands represent 95% confidence intervals. Both stability and species stability were log10 transformed, and species asynchrony was angular transformed.

### Drivers of stability across a latitudinal gradient

A piecewise structural equation model explained 78% of the variation in stability (Fisher’s = 8.24, *P* = 0.61, AICc = 98.24; Fig. 4, Table S8), with species stability and species asynchrony being the main drivers of stability at the small grain (standardized path coefficients = 0.72 and 0.65, respectively). Latitude and soil nutrients influenced stability indirectly. Specifically, latitude increased stability *via* CWM_SLA_, species stability, and species richness (total effect: 0.44, Fig. 4); with CWM_SLA_ affecting stability *via* a negative effect on species stability (standardized path coefficient = −0.14) and stem density *via* a positive effect on species richness (standardized path coefficient = −0.04). Forests with lower soil total nitrogen and soil organic carbon increased stability *via* CWM_SLA_, species stability and stem density (total effect: 0.12, Fig. 4). Negative effects of species richness on stability were both direct (standardized path coefficient = −0.05) and indirect, which operated *via* species stability (standardized path coefficients = −0.17; Fig. 4). In contrast, species richness increased stability *via* species asynchrony (standardized path coefficients = 0.08; Fig. 4).

**Fig. 4.**
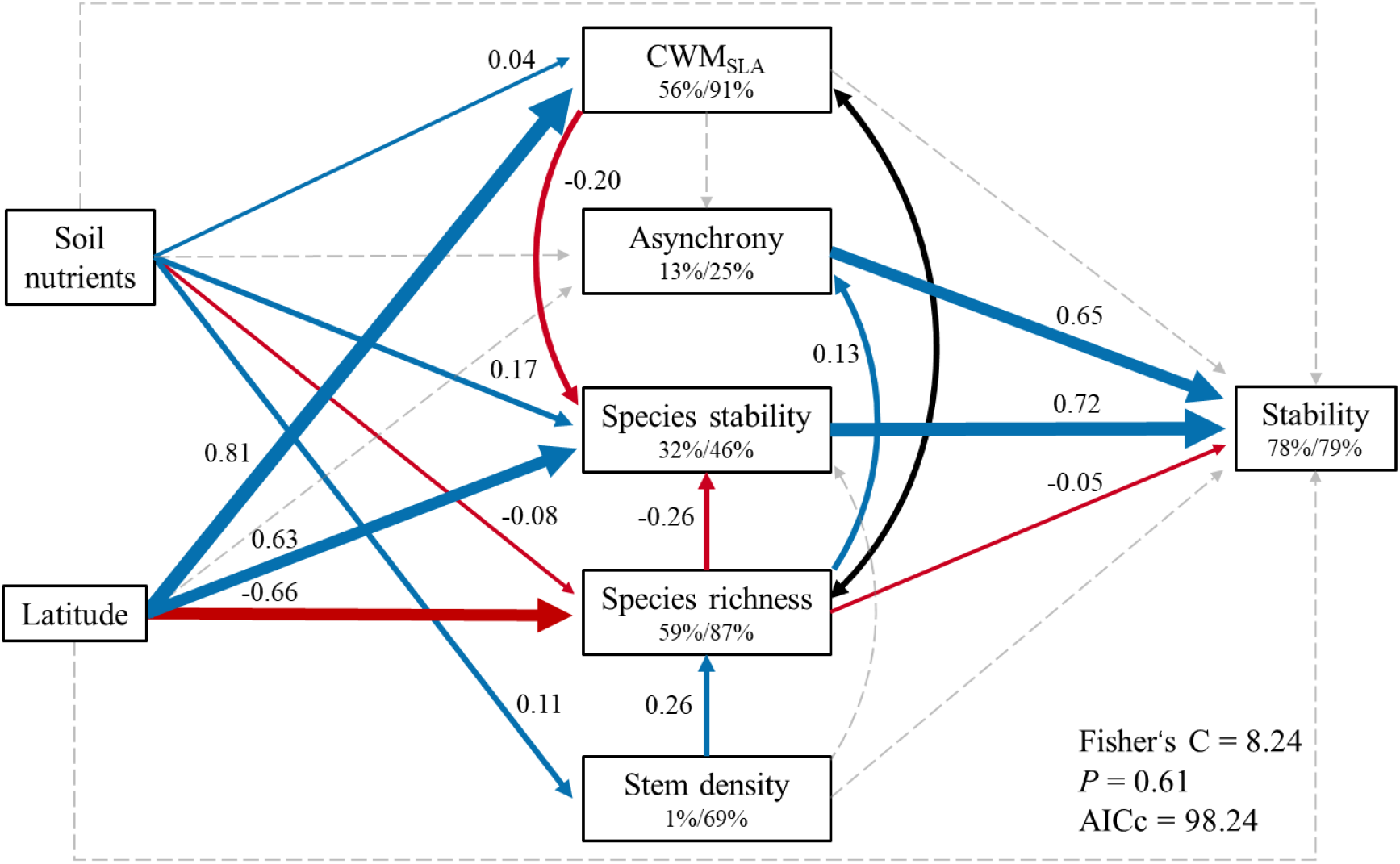
Direct and indirect effects of biotic and abiotic factors on stability across a latitudinal gradient at the small spatial grain. The structural equation model includes soil nutrients (the first principal axis, positive values associated with lower total nitrogen and soil organic carbon), latitude (Latitude), functional trait composition (represented by the community-weighted mean of specific leaf area, CWM_SLA_), species asynchrony (Asynchrony), species stability (Species stability), species richness (Species richness), stem density (Stem density) and stability (Stability). The data fit the model well (Fisher’s C = 8.24, *P* = 0.61; AICc = 98.24). Arrows represent causal relationships between variables. Black bi-directional arrows refer to significant partial pairwise correlations. Solid blue and red lines represent significant (*P* ≤ 0.05) positive and negative standardized paths, respectively. Gray dashed lines represent non-significant paths (*P* > 0.05). Standardized path coefficients are represented for each statistically significant path, and path widths are scaled by standardized path coefficients. Numbers within boxes indicate the variance explained by fixed (left, marginal *R^2^*) and the combination of fixed and random effects (right, conditional *R^2^*).

### Trade-off between species stability and asynchrony in driving stability

We observed a trade-off between the relative importance of species stability and species asynchrony in determining stability (slope = −1.27, *P* = 0.002, *R^2^* = 0.91, Fig. 5). However, this trade-off was not related to latitude. Specifically, there were no significant relationships between latitude and the stabilizing effect of species asynchrony and species stability, and their ratio (*P* value were 0.45, 0.34 and 0.40, respectively. Fig. S6).

**Fig. 5.**
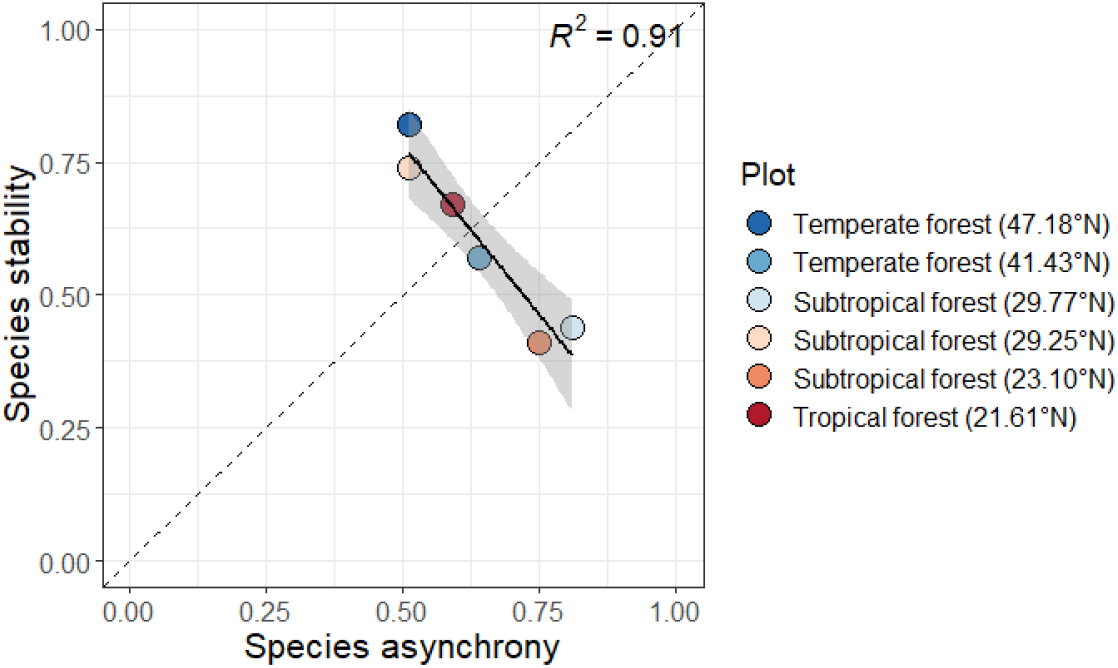
The relative importance of species stability and species asynchrony for stability across a latitudinal gradient. The black line is a linear model fit (*P* < 0.05) for the relationship between the relative importance of both biotic stabilizing mechanisms in determining stability. Each point refers to the direct effect of species stability and asynchrony on stability for each study site.

## Discussion

In naturally-assembled forest ecosystems, stability and its underlying mechanisms vary in response to biotic and abiotic factors^12,14,15^, yet it remains uncertain to what extent these factors shape emergent macroecological patterns of stability across spatial scales. Here, we provide unique insights by demonstrating that there is no clear latitudinal pattern of stability at either small or large spatial grains. Yet, we found that latitude influenced stability *via* biotic stabilizing mechanisms, primarily species stability and species asynchrony, at both spatial grains. Our study therefore reveals consistent positive effects of species stability on stability across spatial grains. The trade-off between species asynchrony and species stability in driving stability did not follow a latitudinal pattern, suggesting that context-dependent factors - to a greater extent than macroecological ones - underlie large-scale patterns of stability in naturally-assembled forests.

Although there was not a clear latitudinal pattern of stability, we found that latitude influences stability *via* biotic stabilizing mechanisms. Latitudinal patterns of species asynchrony and species stability may arise from latitudinal variation in climate and systematic shifts in the importance of abundant and rare species for stability. Warm regions typically hold more species that differ in their response to environmental conditions^41,42^, resulting in higher species asynchrony at lower latitudes as observed in this study. Conversely, the increase in environmental stress (i.e., low temperature) with latitude may lead to a convergence of species responses to the environment, which lowered species asynchrony^43^. One possible explanation for the latitudinal shift in species asynchrony is that it is driven by underlying gradients in species diversity, as suggested by theory that diversity enhance species asynchrony^5,7^, and as observed in this study (Fig. 4). While theory suggests that species asynchrony should increase with spatial grain and extent^16,22^, as both factors are associated with larger species pools, we found that the latitudinal pattern of species asynchrony was weaker at the large spatial grain. For the latitudinal pattern of species stability, shifts in the ecological roles of abundance and rare species may be the primary reason^5^. Our results suggest that the impacts of dominant and rare species on stability change with latitude: while dominant species exhibit strong impacts on stability at higher latitudes with lower community evenness, rare species appear to have stronger impacts on stability at lower latitudes with higher community evenness^5,44^. Furthermore, the contrasting latitudinal patterns for species stability and species richness support growing evidence that species richness has a significantly negative effect on species stability^9,40,44^.

Overall, species richness had neutral or negative effects on stability. This finding suggests that other factors, e.g. climate change, affect biodiversity and stability simultaneously, implying correlation and not causality^28,45^. For example, warming can reduce stability but increase biodiversity, which may result in a neutral or even negative relationship between biodiversity and stability^38,45^. At small spatial grains, our results support the idea that the impacts of species richness on stability are highly context-dependent in naturally-assembled ecosystems^28,40^. On the one hand, species-rich communities increase the likelihood that asynchronous responses among species to environmental conditions increase stability^15,36^. On the other hand, higher functional redundancy in species-rich communities may enhance competition, thereby decreasing stability^28,46^. Therefore, the balance between the diversity of ecological strategies and their redundancy may shape the overall effect of species richness on stability in naturally-assembly forests. At the large spatial grain, the negative effects of species richness on stability are inconsistent with patterns found in grasslands^47^. The destabilizing effects of species richness we observe adds to the growing empirical evidence^18^ that the expected positive effects of macroecological complementarity on ecosystem functioning may not manifest at “small” spatial grains (< 10 ha), a scale at which demographic stochasticity may overwhelm spatial insurance effects^48^.

The trade-off between species stability and asynchrony in driving stability was not driven by latitude, suggesting that the relative importance of different stabilizing mechanisms is not determined uniquely by macroecological drivers. Biotic and abiotic factors operating at smaller spatial scales may shape the relative importance of stabilizing mechanisms. For instance, García-Palacios et al.^30^ found that diversity-stability is mediated by climate and soil nutrients, which were equally as important in stabilizing ecosystem functioning as biotic factors. We found that soil nutrients influenced stability *via* species stability, CWM_SLA_, species richness, and stem density. Soils associated with lower soil organic carbon (SOC), total nitrogen (TN), available phosphorus (AP) and pH had, overall, higher species stability, CWM_SLA_, and stem density, but lower species richness. Yet, when zooming in, the relative importance of soil nutrients on stabilizing mechanisms fluctuated inconsistently across the latitudinal gradient (Supplementary results), likely influencing the overall observed trade-off in the relative importance of species stability and species asynchrony. Therefore, understanding the trade-off between stabilizing mechanisms requires not only the integration of climatic factors and species richness, but also other abiotic conditions and biotic dimensions that operate at different ecological scales^49^.

In conclusion, our study provides fundamental empirical evidence that latitude influences stability in naturally-assembled forest ecosystems *via* stabilizing biotic mechanisms. Furthermore, we show that results from small spatial grains can provide valuable insights for understanding stability and underlying stabilizing mechanisms at spatial extents that are relevant for conservation and forest management. Moreover, our results suggest that ecological processes that alter the functional composition and stem density, like secondary succession, anthropogenic disturbance, or forest management may also play a key role in driving nature’s contributions to people over time.

## Methods

### Study sites and data collection

The dataset used in this study was compiled from CForBio Network (http://www.cfbiodiv.cn), which contains six forest dynamic plots, belonging to three forest ecosystems (temperate forest: LS, CBS; subtropical forest: BDGS, GTS, DHS; tropical forest: BN), whose area ranged from 9 ha to 25 ha (Fig. 1A; Table S1) and that are located across a latitudinal gradient (spanning from 21.61 °N to 47.18 °N). Each plot is divided into subplots (20 m × 20 m = 0.04 ha). Within each plot, all woody plants with diameter at 1.3 m height ≥ 1 cm were measured, mapped, identified, and tagged^50^. All the plots are resurveyed every five years. Our dataset included three inventories for each plot over a period of ten years, yielding a total of more than 1.9 million measurements with 1,013 species.

### Data compilation at the large spatial grain

To explore the variance in the latitudinal pattern of stability and its stabilizing mechanisms among different spatial grains, we compiled a new dataset at the same spatial grain for all study sites (200 m × 200 m = 4.0 ha) including species richness, species asynchrony, species stability and stability, to minimize scale-dependent biases. To this end, we randomly selected 100 subplots with replacement to create an aggregated community. We then calculated species richness, species asynchrony, species stability, and temporal stability (see next section for details). We repeated this process 100 times to obtain a new dataset for each study site (Fig. S1). We also calculated species richness, species asynchrony, species stability and stability using data from all subplots for each plot. From hereon, we refer to results and analyses at the subplot level as the small spatial grain (0.04 ha) and those at the plot level as the large spatial grain (4.0 ha).

### Biotic factors and stability

For each spatial grain at all study sites, we calculated species richness (SR), as the number of living tree species per plot. We then averaged species richness across inventories. Stem density of living trees was also counted and averaged at small spatial grain.

To test the mass ratio hypothesis, we calculated the community-weighted means (CWMs) of functional traits at the small spatial grain using the relative abundance of each species. SLA (specific leaf area, cm^2^/g) was selected to reflect the “fast-slow” strategy of trees^51^, which we calculated with the following formula:

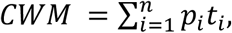

where *p_i_* is the relative abundance of species *i* in a subplot with n species, t_i_ is the mean of SLA at species level for species *i*. SLA of each species was measured by the ratio of leaf area by leaf dry mass.

Species asynchrony at both spatial grains was calculated following ref^6^:

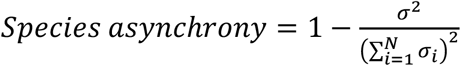

where *σ* is the standard deviation of aboveground biomass (AGB) across three inventories, and *σ_i_* is the standard deviation of AGB of species *i* in each grain with *N* species across the three inventories. A community is perfectly synchronous when the value is equal to 0, and is perfectly asynchronous when the value is equal to 1 (ref^6^). Then, species stability at both spatial grains, i.e., the species level stability weighted by species’ relative abundances, was calculated following ref^5^:

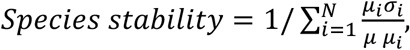

where *μ* and *μ_i_* are the mean of AGB of grain and species *i* within this grain across the three inventories, respectively.

Temporal stability of aboveground biomass (AGB) at both spatial grains was defined as the ratio of mean (*μ*) AGB across three surveys to the standard deviation (*σ*) of AGB^52^. To estimate the aboveground biomass of each individual, we used site- and species-specific allometric equations (see references in Table S2). We then summed aboveground biomass for each inventory.

### Climate and soil nutrients

We obtained monthly temperature, precipitation, and potential evapotranspiration data for each study site from the National Earth System Science Data Center (www.geodata.cn) with a spatial resolution of 1000 m × 1000 m. Given the lag effect of trees’ response to climate, we used climate data for 13 years, including three years before the first inventory at each study site. For all study sites, we calculated mean annual temperature (MAT, °C), mean annual precipitation (MAP, mm^−1^ y^−1^), and mean annual potential evapotranspiration (PET, mm^−1^ y). We also calculated the standard deviation (SD) of MAT, MAP, and PET (Table S3). In addition, we also collected soil nutrient data including soil organic carbon (SOC, g/kg), total nitrogen (TN, mg/g), available phosphorus (AP, mg/kg) and pH within each subplot (Table S3).

We performed principal component analysis (PCA) to reduce the potential collinearity of the climate variables. The first two axes (ClimatePCA1, ClimatePCA2) explained 94% (80% + 14%) of total variation in climatic variables (Table S4; Fig. S2). ClimatePCA1 was positively associated with MAP, MAP, PET and SD of MAP and PET, and negatively associated with SD of MAT. ClimatePCA2 was negatively related to SD of MAP, and positively related to SD of PET. Similarly, we used PCA to reduce the dimensionality of soil nutrient data. The first two PCA axes (SoilPCA1 and SoilPCA2) explained 87% (70% + 17%) of total variation in soil nutrients. SoilPCA1 was negatively associated with SOC, TN, pH and AP, whereas SoilPCA2 was positively and negatively associated with pH and SOC, respectively (Table S4, Fig. S2). As the first principal component for both climate and soil nutrients explained a considerable amount of variation, we used the first principal component axis of both soil nutrients and climate in the following statistical analyses.

### Statistical analyses

Stability and species stability were log10-transformed and species asynchrony was angular transformed (arcsine(square-root(asynchrony))) to meet normality assumptions prior to analysis. First, to test for variation in stability and stabilizing mechanism (species stability and asynchrony) across a latitudinal gradient at both spatial grains, we used linear mixed-effect models with study site as a random intercept at both small and large grains, respectively. Second, to test if there was a variation at spatial grains in bivariate relationships between species richness, species stability, species asynchrony, and stability, we used mixed-effect models with study site as the random effect. Confidence intervals (95%) of mixed effect models were computed using the “ggeffects” package^53^. Diagnostic plots were used to confirm model assumptions, including the homogeneity and normality of model residuals.

To explore the mechanisms shaping stability patterns across the latitudinal gradient at the small spatial grain, we fitted piecewise structural equation models (SEM) across and within study sites^54^. We constructed a hypothetical model based on current theory that we subsequently tested (Fig. S3, Table S5, ref.^14,15,36,38^). This model contains direct paths from functional trait composition (CWM_SLA_), species stability, species richness, species asynchrony, and stem density to stability to assess biotic stabilizing mechanisms, and we included direct paths from soil nutrients (SoilPCA1) and latitude to stability to evaluate abiotic stabilizing mechanisms. We also included paths between biotic mechanisms and between biotic and abiotic mechanisms to represent their indirect effects on stability (Fig. S3, Table S5). We used climate (ClimatePCA1) instead of latitude in an alternative model to test the specific effects of climate on stability (Fig. S4). For the across-plot SEM, we used mixed-effect models with study site as the random effect, and for within study site SEMs, we used generalized least square models. Due to spatial autocorrelation in the response variable (i.e., stability) within study sites (Moran’s I test^55^, *P*-value < 0.05, Table S6), we constructed several variogram models (Exponential, Gaussian spherical, Linear, Rational, Quadratics) to determine the most parsimonious spatial correlation structure. Models with a Gaussian spherical spatial correlation structure had the lowest AIC, and were subsequently used in both across- and within-study site SEMs. For those study sites without strong spatial autocorrelation (LS and CBS), we used generalized least square models without spatial correlation structures in SEM models (Table S6). Directed separation tests were used to examine whether missing paths should be added; we added non-hypothesized, statistically significant path coefficients (*P* < 0.05) to improve model fit. We also removed paths that were not statistically significant to improve model fit. Akaike information criterion (AICc) and Fisher’s C statistic (*P* > 0.05) were used to estimate the goodness-of-fit of model^56^.

To compare the relative importance of the effects of species stability and asynchrony on stability across the latitudinal gradient, we extracted path coefficients from both species stability and asynchrony to stability for each within-plot SEM. We then used a linear model to test whether there is a trade-off in the relative importance of stabilizing mechanisms. Moreover, we also used linear model with path coefficients of species asynchrony and species stability to stability, and their ratio (i.e. asynchrony/species stability) as response variables, and latitude as predictor to test whether this trade-off follow a latitudinal pattern.

All statistical analyses were performed using R v.4.0.3^57^. Moran’s I test and SEM analysis were performed using the “spdpe”^55^ and “piecewiseSEM” packages^54^ respectively, and mixed effects and generalized least squares models were used in the “nlme” package^58^.

## Acknowledgments

We thank the Chinese Forest Biodiversity Monitoring Network (CForBio) for providing the data. We also thank all field technicians and students who have participated in forest inventories at all plots. This work was financially supported by the Strategic Priority Research Program of Chinese Academy of Sciences (XDB31000000), National Natural Science Foundation of China (32171536, 32061123003 and 31670441). Nathaly R. Guerrero-Ramírez thanks the Dorothea Schlözer Postdoctoral Programme of the Georg-August-Universität for their support, and Dylan Craven acknowledges funding from FONDECYT Project 1201347.

## Authors’ Contributions

X.J.Q. and M.X.J. designed the research; T.Y.Z., N.R.G.-R., D.C. and X.J.Q. conceived ideas; T.Y.Z., compiled and analyzed the data with the help of N.R.G.-R., D.C., H.K. and X.J.Q.; T.Y.Z., N.R.G.-R., D.C., H.K. and X.J.Q. led the writing of the manuscript. All authors revised the drafts and gave final approval for publication.

## Competing interests

The authors declare no competing interests.

## Notes

### Competing Interest Statement

The authors have declared no competing interest.

