## supplemental Table and Figure for "Latitude influences stability via stabilizing mechanisms in naturally-assembled forest ecosystems at different spatial grains"

### Supplementary information

### Supplementary results

#### Climatic variation underlying latitudinal patterns

The SEM model testing drivers of stability across the latitudinal gradient indicated that species stability, species asynchrony, and climates were the key drivers of stability at the small grain. Overall, the SEM explained 78% of total variation in stability, which increased to 79% when accounting for random effects. Overall, our model fit the data well (Fisher's = 8.49, *P* = 0.39, AICc = 124.49, Table S9). Specifically, both species stability and asynchrony promoted stability directly (standardized path coefficients = 0.72 and 0.65). Climates enhanced stability indirectly via $CWM_{SLA}$, species stability and species richness (total effect: -0.47). Unexpectedly, species richness had a direct, negative effect on stability (standardized path coefficient = -0.05), and also destabilized AGB via species stability. However, it can also improve stability via a positive effect on species asynchrony (Figure S4, Table S9). Although $CWM_{SLA}$ and stem density had no direct effect on stability, they affected stability indirectly via a negative effect on species stability and a positive effect on biodiversity, respectively. Our finding showed that soil nutrients had a minor effect on stability directly (standardized path coefficient = -0.02), but indirectly affects stability via biotic factors, including a positive effect on $CWM_{SLA}$, species stability and stem density and a negative effect on species richness (total effect: 0.12, Figure S4)

#### Plot level drivers of stability

When considering study site separately, SEMs explained between 59% and 93% of variation in stability at the small spatial grain and confirmed that both species stability and asynchrony are key drivers of temporal stability (Fig. S5, Table S10-S15). For five of the six study sites, we found that species richness had a neutral, direct effect on stability (Fig. S5). In contrast, we found an indirect, negative effect of species richness on stability *via* species stability for four of the six study sites (Fig. S5), while species richness only stabilized AGB *via* species asynchrony in one of the six study sites (Fig. S5). Similarly, $CWM_{SLA}$ played a minor role in stabilizing AGB, reducing stability *via* species stability in three of the six study sites. Although stem density had a positive effect on species richness, it had a minor effect on stability. Also, soil nutrients affected stability indirectly *via* biotic factors, whose effect varies across study sites (Fig. S5).

### Supplementary figures

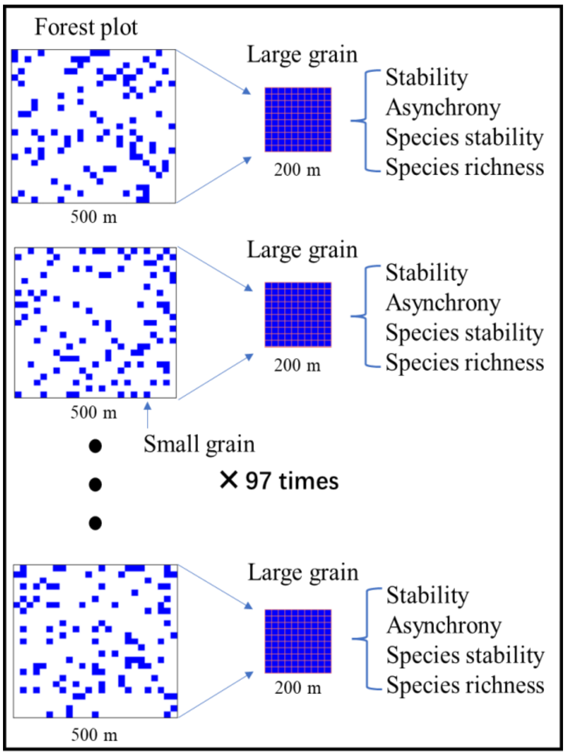


**Fig. S1 Sketch showing the process for conducting a community at large grain (200 m × 200 m).** We randomly selected 100 subplots (20 m, small grain) with replacement to create an aggregated community for each plot. Then, we calculated the species richness, asynchrony, species stability and community stability at this grain according to the materials and methods section. We repeated this process 100 times.

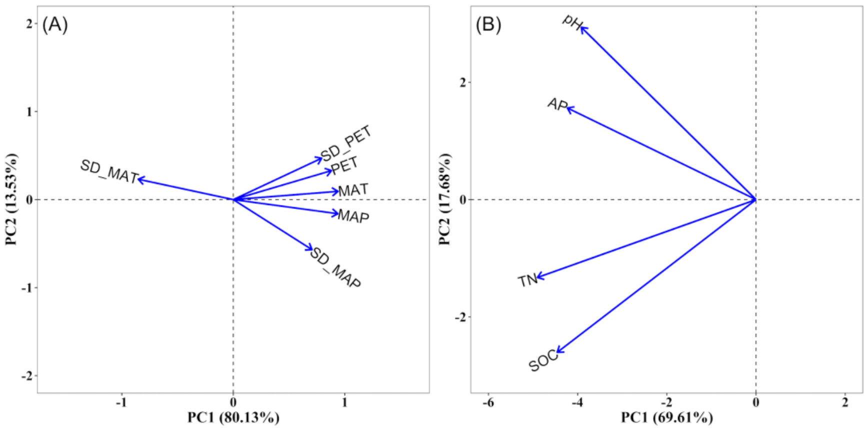


**Fig. S2 Principal component analysis for climate factors (A) and soil nutrients (B).** Climatic factors included: mean annual temperature (MAT, °C; PC1: 0.94, PC2: 0.09), mean annual precipitation (MAP, mm; PC1: 0.94, PC2: -0.16) and mean annual potential evapotranspiration (PET, mm; PC1: 0.88, PC2: 0.33), and the standard deviation (SD) of MAT (SD_MAT; PC1: -0.85, PC2: 0.23), MAP (SD_MAP; PC1: 0.71, PC2: -0.57) and PET (SD_PET; PC1: 0.79, PC2: 0.47). Soil nutrients included: soil organic carbon (SOC, g/kg; PC1: -4.46, PC2: -2.60), total nitrogen (TN, mg/g; PC1: -4.90, PC2: -1.33), available phosphorus (AP, mg/kg; PC1: -4.23, PC2: 1.56) and pH (pH; PC1: -3.92, PC2: 2.94).

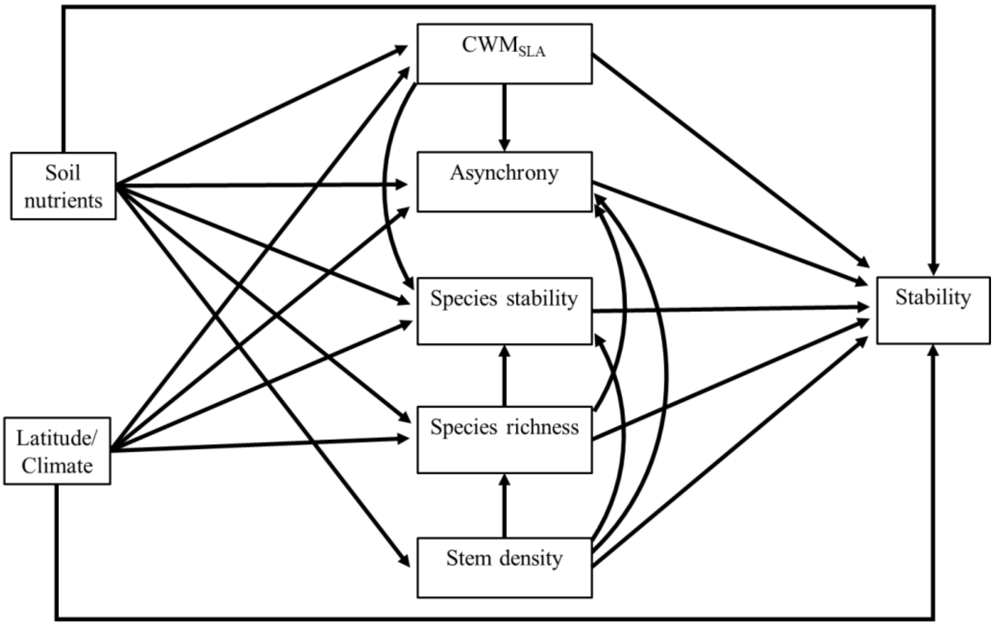


**Fig. S3 The conceptual model linking community stability with the effects of abiotic and biotic factors.** The arrows indicate the causal relationship, theoretical support for potential paths in Table S5. The structural equation model included soil nutrients (the first axis of PCA for soil nutrients, Soil nutrients), latitude or Climate (Latitude/Climate), functional trait composition (SLA, $CWM_{SLA}$), species asynchrony (Asynchrony), species stability (Species stability), species richness (Species richness), stem density (Stem density) and community temporal stability (Stability).

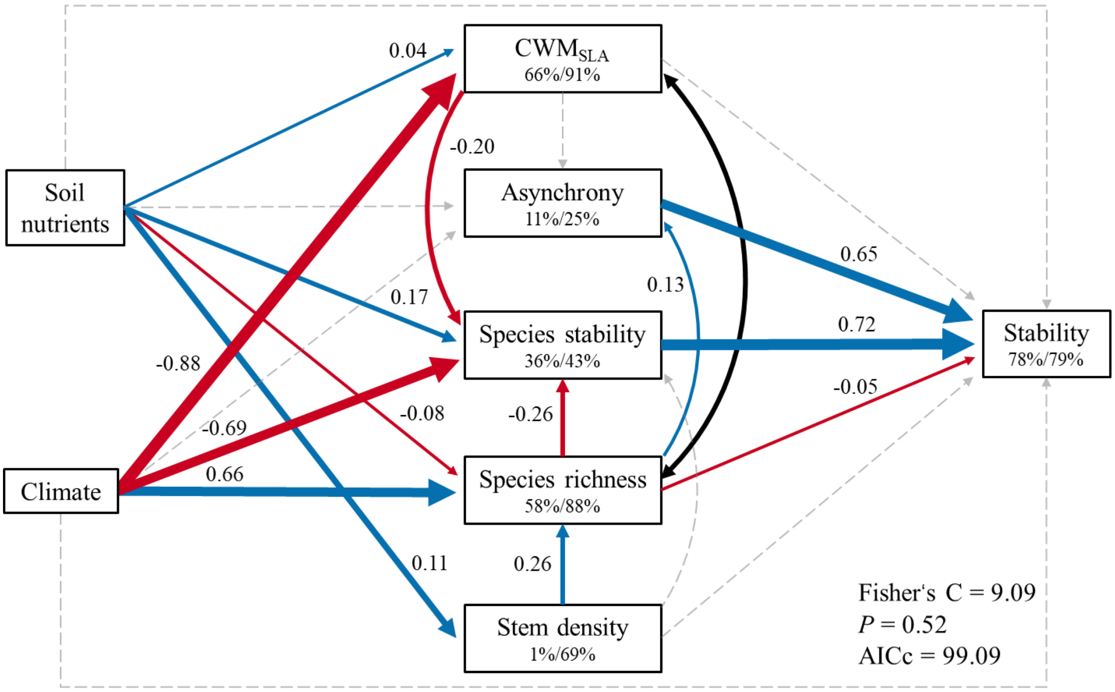


**Fig. S4 Direct and indirect effect of biotic and abiotic factors on stability across a latitudinal spatial extent at small spatial grain.** The structural equation model included soil nutrients (the first principal axis, positive values associated with lower total nitrogen and soil carbon), climate factors (the first principal axis, positive values associated with higher MAP and MAT, negative values associated with lower standard deviation of MAT), functional trait composition (represented by the community-weighted mean of specific leaf area, $CWM_{SLA}$), species asynchrony (Asynchrony), species stability (Species stability), species richness (Species richness), stem density (Stem density) and stability (Stability). The data fit the model well (Fisher's C = 9.09, $P = 0.52$; AICc = 99.09). Arrows represent causal relationships between variables. Black bi-directional arrows refer to significant partial pairwise correlations. Solid blue and red lines represent significant ($P \leq 0.05$) positive and negative standardized paths, respectively. Grey dashed lines represent non-significant paths ($P > 0.05$). Standardized path coefficients are represented to each (significant) path, and widths of significant paths are scaled by standardized path coefficients. These numbers within the boxes indicate the variation explained by fixed (left, marginal $R^2$) and combination of fixed and random effects (right, conditional $R^2$).

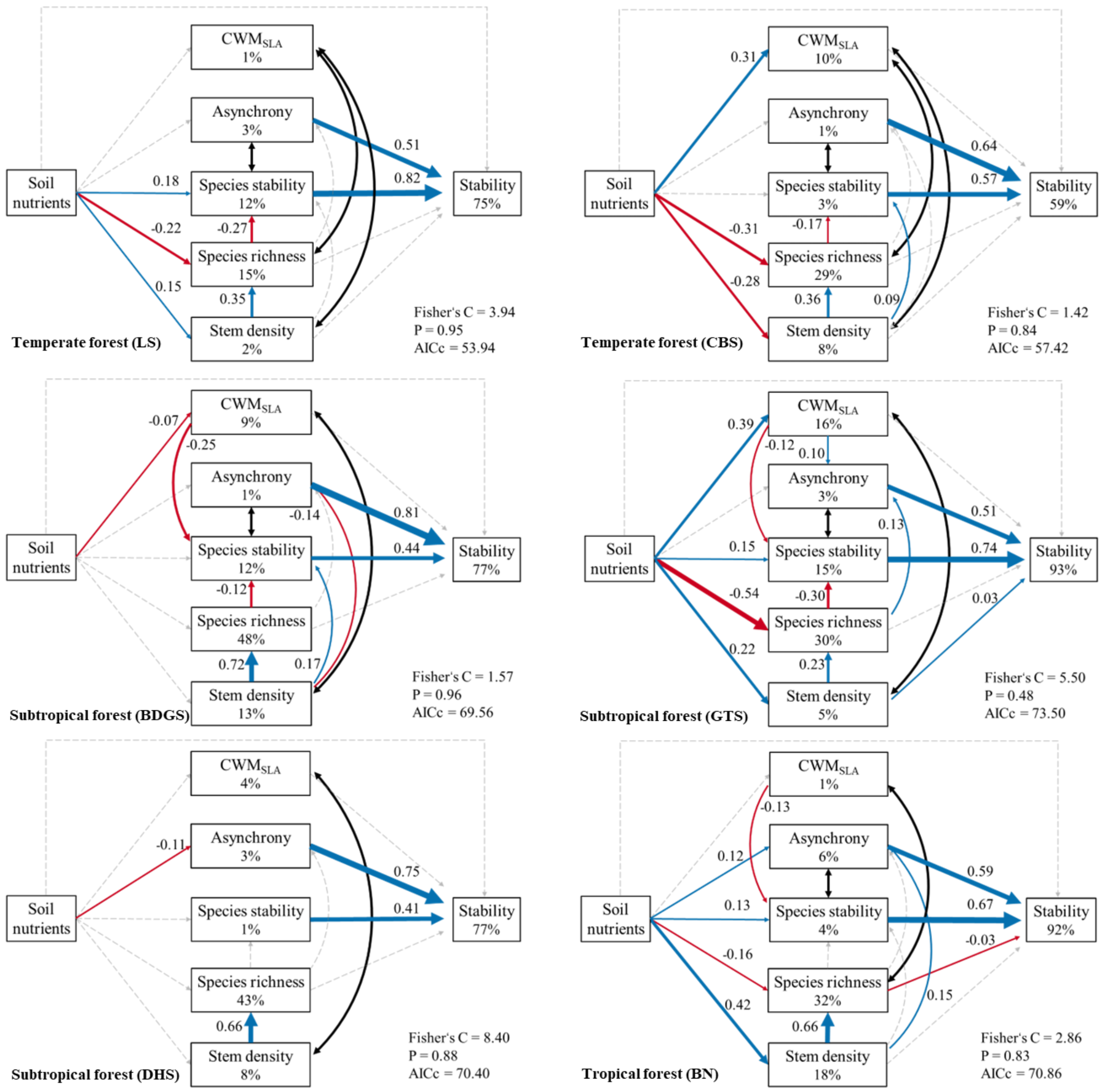


**Fig. S5 Direct and indirect effects of biotic and abiotic factors on stability for each study site.** These structural equation models included soil nutrients (the first principal axis, positive values associated with lower total nitrogen and soil organic carbon), functional trait composition (represented by the community-weighted mean of specific leaf area, $CWM_{SLA}$), species asynchrony (Asynchrony), species stability (Species stability), species richness (Species richness), stem density (Stem density) and stability (Stability). Arrows represent causal relationships between variables. Black bi-directional arrows refer to significant partial correlations. Solid blue and red lines represent significant ($P \leq 0.05$) positive and negative standardized paths, respectively, and grey dashed lines represent non-significant paths. Standardized path coefficients represent each statistically significant path, and widths of paths are scaled by standardized path coefficients. The variation in $CWM_{SLA}$, asynchrony, species stability, species richness, stem density and stability explained by influence is represented in these boxes. The study site name is indicated in the lower left corner of each figure, and panels are ordered from top to bottom according to the latitude of the study sites from north to south.

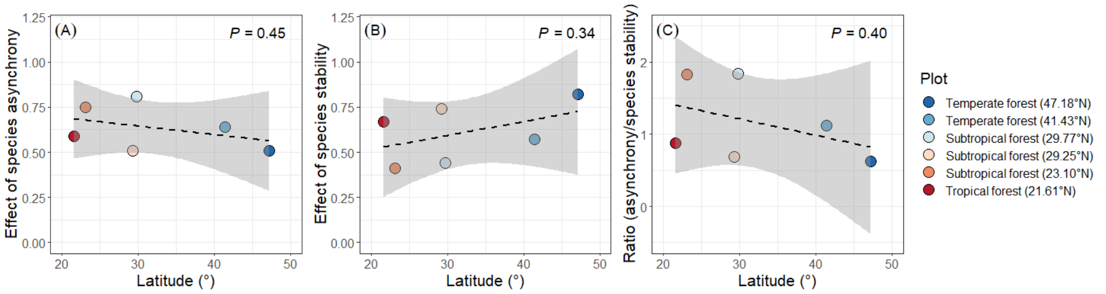


**Fig. S6 Latitudinal pattern of stabilizing effect of species asynchrony (A) and species stability (B), and their ratio (species asynchrony/species stability) (C).** The dashed black line is a linear model fit ($P > 0.05$) for the relationship between latitude and stabilizing effect of species asynchrony and species stability, and their ratio. Each point refers to the direct effect of species stability and asynchrony on stability for each study site.

Table S1 General description of each study site.

| Plot | Plot Abbreviation | Forest type | Area (ha) | No. of quadrat (20m × 20 m) | Latitude (°) | Longitude (°) | No. of species | Survey year | | |
|---|---|---|---|---|---|---|---|---|---|---|
| | | | | | | | | 1st | 2nd | 3rd |
| Liangshui | LS | Temperate | 9 | 225 | 47.18 | 128.88 | 44 | 2006 | 2011 | 2016 |
| Changbaishan | CBS | Temperate | 25 | 625 | 41.43 | 127.42 | 52 | 2004 | 2009 | 2014 |
| Badagongshan | BDGS | Subtropical | 25 | 625 | 29.77 | 110.09 | 232 | 2011 | 2016 | 2021 |
| Gutianshan | GTS | Subtropical | 24 | 600 | 29.25 | 118.12 | 159 | 2005 | 2010 | 2015 |
| Dinghushan | DHS | Subtropical | 20 | 500 | 23.10 | 112.32 | 210 | 2005 | 2010 | 2015 |
| Banna | BN | Tropical | 20 | 500 | 21.61 | 101.57 | 468 | 2007 | 2012 | 2017 |

Table S2 The references of biomass equations for each study site

| Plot | References |
|---|---|
| LS | Wang, 2006; Li et al., 2010; Zhou et al., 2018 |
| CBS | Wang, 2006; Chen & Zhu. 1989; Li et al., 2010; Zhou et al., 2018 |
| BDGS | Xu, 2016; Zhou et al., 2018 |
| GTS | Lin et al., 2015; Zhang et al., 2007; Zhou et al., 2018 |
| DHS | Chen et al., 2015; Zhou et al., 2018 |
| BN | Lv et al., 2007; Linger et al., 2020; Zhou et al., 2018 |

Table S3 The basic information about abiotic factors.

| Abiotic factors | | LS | CBS | BDGS | GTS | DHS | BN |
|---|---|---|---|---|---|---|---|
| Climate | MAT (°C) | 2.1 | 3.95 | 11.91 | 16.25 | 20.93 | 21.87 |
| | MAP (mm) | 511.1 | 588.0 | 1260.3 | 1478.2 | 1472.8 | 1447.7 |
| | PET (mm/year) | 645.2 | 681.2 | 795.9 | 882.8 | 1042.5 | 1152.3 |
| | SD of MAT | 0.65 | 0.62 | 0.30 | 0.33 | 0.38 | 0.35 |
| | SD of MAP | 83.58 | 99.36 | 152.18 | 309.38 | 229.08 | 154.76 |
| | SD of PET | 20.10 | 24.20 | 24.80 | 28.30 | 27.40 | 37.10 |
| Soil | Soil organic carbon (g/mg) | 75.70 | 95.30 | 83.10 | 43.60 | 35.40 | 18.40 |
| | Total nitrogen (g/mg) | 8.40 | 6.38 | 6.19 | 2.22 | 1.18 | 1.83 |
| | Available Phosphorus (mg/kg) | 8.83 | 8.49 | 6.34 | 1.23 | 1.80 | 4.80 |
| | PH | 5.77 | 5.45 | 4.58 | 4.63 | 3.75 | 4.92 |

Mean and standard variation value of climate factors across 13 years for each study site, and mean value of soil nutrients for each quadrat are shown in this table. MAT: mean annual temperature; MAP: mean annual precipitation; PET: mean annual potential evapotranspiration; SD_MAT: Standard deviation of MAT; SD_MAP: Standard deviation of MAP; SD_PET: Standard deviation of PET.

Table S4 The PCA analysis for climate factors and soil nutrients

| Abiotic factors | | PCA1 | PCA2 |
|---|---|---|---|
| Climate | MAT | 0.94 | 0.09 |
| | SD of MAT | -0.85 | 0.23 |
| | MAP | 0.94 | -0.16 |
| | SD of MAP | 0.71 | -0.57 |
| | PET | 0.88 | 0.33 |
| | SD of PET | 0.79 | 0.47 |
| Eigenvalue | | 4.81 | 0.81 |
| Cumulative Proportion (%) | | 0.80 | 0.94 |
| Soil | SOC | -4.46 | -2.60 |
| | TN | -4.90 | -1.33 |
| | AP | -4.23 | 1.56 |
| | pH | -3.92 | 2.94 |
| Eigenvalue | | 2.78 | 0.71 |
| Cumulative Proportion (%) | | 0.70 | 0.87 |

SOC: soil organic carbon; TN: total nitrogen; AP: available phosphorus; PH: soil pH.

Table S5 Illustration of ecological paths and theories

| Hypothesized pathway | | Ecological theories | References |
|---|---|---|---|
| predictor | response | | |
| Soil nutrients | Stability | Changes in soil nutrients could mediate species composition, community structure and biodiversity that also resulted in the variation of community stability consequently. | Morin et al., 2014; Grman et al., 2010 |
| Soil nutrients | $CWM_{SLA}$ | Plant modulates its functional traits to the specific environmental condition due to traits plasticity. | Morin et al., 2014 |
| Soil nutrients | Species stability | Soil nutrients affect plant growth and the interaction between species. | Ma et al., 2017 |
| Soil nutrients | Species richness | Soil nutrients increase the niche and thus biodiversity, or decrease biodiversity by enhancing the competition among species. | Chen et al., 2020; Grman et al., 2010 |
| Soil nutrients | Asynchrony | Soil nutrients affect asynchrony by modifying the competition among species, species turnover. | Ma et al., 2017 |
| Soil nutrients | Stem density | Fertile soil providing better environmental conditions can afford more individuals. | Ouyang et al., 2019 |
| Latitude/Climate | Stability | Climate directly affects plant growth, community structure and composition, and ecosystem functions, i.e., productivity and stability. | Ma et al., 2017; Chen et al., 2021; García-Palacios et al., 2018 |
| Latitude/Climate | $CWM_{SLA}$ | Plants improve fitness for survival by regulating traits under specific climatic conditions. For example, under drought conditions, the reduced leaf area reduces loss. | García-Palacios et al., 2018 |
| Latitude/Climate | Species stability | Climate directly affects growth, mortality of species and thus stability. | Ma et al., 2017 |
| Latitude/Climate | Species richness | Climate is the key driver of biodiversity at the regional scale, as depicted by the latitudinal diversity gradient with species richness increase from the poles to the equator. | Chu et al., 2018; Gaston, 2008 |
| Latitude/Climate | Asynchrony | Climate impacts species asynchrony indirectly via changing community composition. | Usinowicz et al., 2017; Ma et al., 2017 |
| $CWM_{SLA}$ | Stability | Functional traits can drive community stability effectively. For example, communities dominated by conservative traits (e.g., higher leaf carbon content, leaf thickness, leaf dry matter content) preferred stable than that by explorative traits (e.g., higher leaf phosphorus, leaf area, specific leaf area) under environmental disturbance. | Craven et al., 2018; Schnabel et al., 2021 |

| | | | |
|---|---|---|---|
| $CWM_{SLA}$ | Species stability | Functional traits can reflect the growth strategy of species population and the species stability | Schnabel et al., 2021; García-Palacios et al., 2018; Majeková et al., 2014 |
| $CWM_{SLA}$ | Asynchrony | Functional traits affect plant growth and thus community dynamics. generally, dominant species with conservative traits often fluctuate less, while those with explorative traits have higher temporal variability. | Dolezal et al., 2020 |
| Species stability | Stability | Species stability is the key driver of community stability due to the new ecological frame. | Thibaut & Connolly, 2013; Xu et al., 2021 |
| Species richness | Species stability | The increase of species with similar ecological functions will increase the competition between species and thus reduce species stability. | Hector et al., 2010; Xu et al., 2021 |
| Species richness | Asynchrony | Due to the unique response to environmental disturbance of each species, increased species richness could enhance community asynchrony. | Loreau, M. & de Mazancourt, 2008; Craven et al., 2018 |
| Species richness | Stability | Species mixtures with a higher potential capacity to buffer against environmental fluctuation than monoculture and thus higher stability benefiting from the facilitation or competition reduction among species. | McCann et al., 2000; Bai et al., 2004; van der Plas 2019 |
| Asynchrony | Stability | Asynchrony refers to the inconsistent response to environmental disturbance, which is regarded as the key driver of community stability. | Loreau, M. & de Mazancourt, 2008; Craven et al., 2018 |
| Stem density | Species stability | Stem density can mediate the interaction among individuals, thus the species stability. | Jucker et al., 2014 |
| Stem density | Species richness | More individuals could represent more species because of a statistical effect. | Chu et al., 2018 |
| Stem density | Asynchrony | Stem density enhances the species asynchrony potentially due to communities with higher stem density having a higher probability to contain more species. | Dolezal et al., 2020 |
| Stem density | Stability | Stem density can mediate the interaction among individuals, thus the community stability. | Jucker et al., 2014; Del Rio et al., 2017 |

Table S6 The results of spatial autocorrelation for stability of each study site

| Plot | Moran's I | | Expectation | Variance | *P* |
|---|---|---|---|---|---|
| | Observed | Standard deviation | | | |
| LS | 0.03 | 0.79 | -0.004 | 0.002 | 0.214 |
| CBS | 0.04 | 1.42 | -0.001 | 0.000 | 0.078 |
| GTS | 0.14 | 4.72 | -0.002 | 0.001 | 0.001 |
| BDGS | 0.07 | 2.62 | -0.001 | 0.001 | 0.004 |
| DHS | 0.08 | 2.44 | -0.002 | 0.001 | 0.007 |
| BN | 0.17 | 5.28 | -0.002 | 0.001 | 0.001 |

Expectation: its expectation under the method assumption; Variance: its variance under the method assumption; *P*: P value.

Table S7 Mixed effect models testing the effect of species richness, species asynchrony and species stability on stability.

| Spatial grain | Fixed Effects | AICc | t | *P*-value | $R^2m$ | $R^2c$ |
|---|---|---|---|---|---|---|
| Small | Species richness | 1922.41 | -4.26 | <0.001 | 0.03 | 0.10 |
| | Asynchrony | 489.77 | 42.97 | <0.001 | 0.32 | 0.57 |
| | species stability | 557.44 | 41.67 | <0.001 | 0.45 | 0.49 |
| Large | Species richness | -710.27 | -3.67 | <0.001 | 0.53 | 0.84 |
| | Asynchrony | -1443.41 | 38.11 | <0.001 | 0.30 | 0.97 |
| | Species stability | -826.42 | 11.3 | <0.001 | 0.67 | 0.92 |

These bivariate relationships were explored at both small and large spatial scales. AICc is the Akaike's information criterion; t is the ratio between the estimate and its standard error; *P*-value from a t-distribution; $R^2m$ and $R^2c$ refer to the variance explained by fixed effects and the combination of fixed and random effects.

Table S8 The standardized path coefficients of structural equation models testing the impact of biotic and abiotic factors (Latitude) on community stability for across-plot analysis. Models are presented in Fig. 4. Significant effects (*P* < 0.05) are indicated in bold.

| Response | Predictor | Crit. Value | Non-standardized estimate | Standardized estimate | *P*-value |
|---|---|---|---|---|---|
| Stability | Species richness | -2.10 | 0.00 | -0.05 | **0.036** |
| Stability | Asynchrony | 68.56 | 0.70 | 0.65 | **0.000** |
| Stability | Soil nutrients | -1.14 | -0.04 | -0.02 | 0.254 |
| Stability | Latitude | -0.14 | 0.00 | -0.01 | 0.895 |
| Stability | Species stability | 65.88 | 0.93 | 0.72 | **0.000** |
| Stability | Stem density | 1.70 | 0.00 | 0.03 | 0.090 |
| Stability | $CWM_{SLA}$ | 1.72 | 0.00 | 0.04 | 0.085 |
| Species stability | Soil nutrients | 4.47 | 0.25 | 0.17 | **0.000** |
| Species stability | Latitude | 4.07 | 0.02 | 0.63 | **0.015** |
| Species stability | Species richness | -6.32 | -0.01 | -0.26 | **0.000** |
| Species stability | $CWM_{SLA}$ | -4.47 | 0.00 | -0.20 | **0.000** |
| Species stability | Stem density | 1.07 | 0.00 | 0.03 | 0.286 |
| Asynchrony | Species richness | 2.92 | 0.00 | 0.13 | **0.004** |
| Asynchrony | Soil nutrients | 1.44 | 0.10 | 0.06 | 0.151 |
| Asynchrony | Latitude | -1.49 | -0.01 | -0.21 | 0.210 |
| Species richness | Soil nutrients | -4.46 | -6.65 | -0.08 | **0.000** |
| Species richness | Stem density | 21.17 | 0.03 | 0.26 | **0.000** |
| Species richness | Latitude | -3.45 | -1.18 | -0.66 | **0.026** |
| Stem density | Soil nutrients | 3.95 | 82.43 | 0.11 | **0.000** |
| $CWM_{SLA}$ | Soil nutrients | 2.47 | 12.84 | 0.04 | **0.014** |
| $CWM_{SLA}$ | Latitude | 3.47 | 5.54 | 0.81 | **0.026** |
| Partial correlations | | | | | |
| Species richness | $CWM_{SLA}$ | -6.05 | -0.11 | -0.11 | **0.000** |

Table S9 The standardized estimates of structural equation models testing the impact of biotic and abiotic factors (Climate) on community stability for across-plot analysis. Models are presented in Figure S4. Significant effects ($P$ < 0.05) are indicated in bold.

| Response | Predictor | Crit. Value | Non-standardized estimate | Standardized estimate | *P*-value |
|---|---|---|---|---|---|
| Stability | Species richness | -2.11 | 0.00 | -0.05 | **0.035** |
| Stability | Asynchrony | 68.58 | 0.70 | 0.65 | **0.000** |
| Stability | Soil nutrients | -1.15 | -0.05 | -0.02 | 0.250 |
| Stability | Climate | 0.18 | 0.00 | -0.01 | 0.868 |
| Stability | Species stability | 65.79 | 0.93 | 0.72 | **0.000** |
| Stability | Stem density | 1.69 | 0.00 | 0.03 | 0.091 |
| Stability | $CWM_{SLA}$ | 1.73 | 0.00 | 0.04 | 0.084 |
| Species stability | Soil nutrients | 4.47 | 0.25 | 0.17 | **0.000** |
| Species stability | Climate | -6.12 | -0.22 | -0.69 | **0.004** |
| Species stability | Species richness | -6.20 | 0.00 | -0.26 | **0.000** |
| Species stability | $CWM_{SLA}$ | -4.51 | 0.00 | -0.20 | **0.000** |
| Species stability | Stem density | 1.09 | 0.00 | 0.03 | 0.278 |
| Asynchrony | Species richness | 2.98 | 0.00 | 0.13 | **0.003** |
| Asynchrony | Soil nutrients | 1.47 | 0.11 | 0.06 | 0.142 |
| Asynchrony | Climate | 1.19 | 0.07 | 0.18 | 0.301 |
| Species richness | Soil nutrients | -4.45 | -6.65 | -0.08 | **0.000** |
| Species richness | Stem density | 21.15 | 0.03 | 0.26 | **0.000** |
| Species richness | Climate | 3.19 | 11.44 | -0.66 | **0.033** |
| Stem density | Soil nutrients | 3.95 | 82.43 | 0.11 | **0.000** |
| $CWM_{SLA}$ | Soil nutrients | 2.47 | 12.88 | 0.04 | **0.013** |
| $CWM_{SLA}$ | Climate | -4.58 | -58.09 | -0.88 | **0.010** |
| Partial bivariate correlations | | | | | |
| Species richness | $CWM_{SLA}$ | -6.05 | -0.11 | -0.11 | **0.000** |

Table S10 The standardized estimates of structural equation models testing the impact of biotic and abiotic factors on community stability for LS plot. Models are presented in Fig. S5. Significant effects ($P$ < 0.05) are indicated in bold.

| Response | Predictor | Crit. Value | Non-standardized estimate | Standardized estimate | *P* -value |
|---|---|---|---|---|---|
| Stability | Soil nutrients | 0.71 | 0.06 | 0.03 | 0.480 |
| Stability | Asynchrony | 14.58 | 0.43 | 0.51 | **0.000** |
| Stability | Species stability | 21.96 | 0.98 | 0.81 | **0.000** |
| Stability | Species richness | 1.29 | 0.01 | 0.05 | 0.197 |
| Stability | Stem density | -1.88 | 0.00 | -0.07 | 0.061 |
| Asynchrony | Soil nutrients | -1.63 | -0.32 | -0.11 | 0.106 |
| Asynchrony | Species richness | 1.96 | 0.02 | 0.13 | 0.051 |
| Species stability | Soil nutrients | 2.75 | 0.36 | 0.18 | **0.006** |
| Species stability | Species richness | -4.24 | -0.03 | -0.27 | **0.000** |
| $CWM_{SLA}$ | Soil nutrients | 1.67 | 25.51 | 0.11 | 0.097 |
| Species richness | Soil nutrients | -3.49 | -3.51 | -0.22 | **0.001** |
| Species richness | Stem density | 5.52 | 0.03 | 0.35 | **0.000** |
| Stem density | Soil nutrients | 2.20 | 24.89 | 0.15 | **0.029** |
| Partial bivariate correlations | | | | | |
| Species stability | Asynchrony | -2.33 | -0.15 | -0.15 | **0.010** |
| Stem density | $CWM_{SLA}$ | 3.65 | 0.24 | 0.24 | **0.000** |
| Species richness | $CWM_{SLA}$ | -2.88 | -0.19 | -0.19 | **0.002** |

Table S11 The standardized estimates of structural equation models testing the impact of biotic and abiotic factors on community stability for CBS plot. Models are presented in Fig. S5. Significant effects ($P$ < 0.05) are indicated in bold.

| Response | Predictor | Crit. Value | Non-standardized estimate | Standardized estimate | *P*-value |
|---|---|---|---|---|---|
| Stability | Soil nutrients | -1.34 | -0.13 | -0.04 | 0.182 |
| Stability | Asynchrony | 24.33 | 0.56 | 0.64 | **0.000** |
| Stability | Species stability | 21.28 | 0.89 | 0.57 | **0.000** |
| Stability | Species richness | -0.94 | 0.00 | -0.03 | 0.346 |
| Stability | Stem density | 1.18 | 0.00 | 0.03 | 0.238 |
| Stability | $CWM_{SLA}$ | -0.98 | 0.00 | -0.03 | 0.328 |
| Asynchrony | Species richness | 1.78 | 0.01 | 0.08 | 0.076 |
| Asynchrony | Soil nutrients | 1.10 | 0.19 | 0.05 | 0.273 |
| Asynchrony | Stem density | -1.12 | 0.00 | -0.05 | 0.264 |
| Species stability | Species richness | -3.57 | -0.02 | -0.17 | **0.000** |
| Species stability | Soil nutrients | 0.65 | 0.06 | 0.03 | 0.517 |
| Species stability | Stem density | 2.10 | 0.00 | 0.09 | **0.036** |
| $CWM_{SLA}$ | Soil nutrients | 8.22 | 56.58 | 0.31 | **0.000** |
| Species richness | Soil nutrients | -8.83 | -7.10 | -0.31 | **0.000** |
| Species richness | Stem density | 10.18 | 0.06 | 0.36 | **0.000** |
| Stem density | Soil nutrients | -7.19 | -37.30 | -0.28 | **0.000** |
| Partial bivariate correlations | | | | | |
| Stem density | $CWM_{SLA}$ | -3.24 | -0.13 | -0.13 | **0.001** |
| Species richness | $CWM_{SLA}$ | 3.09 | 0.12 | 0.12 | **0.001** |
| Species stability | Asynchrony | -4.93 | -0.19 | -0.19 | **0.000** |

Table S12 The standardized estimates of structural equation models testing the impact of biotic and abiotic factors on community stability for BDGS plot. Models are presented in Fig. S5. Significant effects ($P$ < 0.05) are indicated in bold.

| Response | Predictor | Crit. Value | Non-standardized estimate | Standardized estimate | *P*-value |
|---|---|---|---|---|---|
| Stability | Soil nutrients | 0.51 | 0.03 | 0.01 | 0.607 |
| Stability | Asynchrony | 41.19 | 0.89 | 0.81 | **0.000** |
| Stability | Species stability | 21.09 | 0.87 | 0.44 | **0.000** |
| Stability | Species richness | -0.35 | 0.00 | -0.01 | 0.725 |
| Stability | $CWM_{SLA}$ | 1.38 | 0.00 | 0.03 | 0.167 |
| Asynchrony | Soil nutrients | 0.11 | 0.01 | 0.00 | 0.910 |
| Asynchrony | Species richness | 0.96 | 0.00 | 0.05 | 0.335 |
| Asynchrony | Stem density | -2.66 | 0.00 | -0.15 | **0.008** |
| Species stability | Soil nutrients | -0.20 | -0.01 | -0.01 | 0.838 |
| Species stability | Species richness | -2.24 | 0.00 | -0.12 | **0.026** |
| Species stability | Stem density | 2.72 | 0.00 | 0.17 | **0.007** |
| Species stability | $CWM_{SLA}$ | -4.89 | 0.00 | -0.25 | **0.000** |
| Species richness | Soil nutrients | -0.94 | -2.30 | -0.03 | 0.348 |
| Species richness | Stem density | 21.74 | 0.05 | 0.72 | **0.000** |
| Stem density | Soil nutrients | 1.05 | 40.51 | 0.04 | 0.292 |
| $CWM_{SLA}$ | Soil nutrients | -2.23 | -20.67 | -0.07 | **0.026** |
| Partial bivariate correlations | | | | | |
| Stem density | $CWM_{SLA}$ | -17.38 | -0.57 | -0.57 | **0.000** |
| Species stability | Asynchrony | -2.24 | -0.09 | -0.09 | **0.013** |

Table S13 The standardized estimates of structural equation models testing the impact of biotic and abiotic factors on community stability for GTS plot. Models are presented in Fig. S5. Significant effects ($P$ < 0.05) are indicated in bold.

| Response | Predictor | Crit. Value | Non-standardized estimate | Standardized estimate | *P*-value |
|---|---|---|---|---|---|
| Stability | Species richness | -1.52 | 0.00 | -0.02 | 0.128 |
| Stability | Asynchrony | 46.73 | 0.70 | 0.51 | **0.000** |
| Stability | Species stability | 62.02 | 0.95 | 0.74 | **0.000** |
| Stability | Soil nutrients | 1.38 | 0.11 | 0.02 | 0.169 |
| Stability | $CWM_{SLA}$ | 1.70 | 0.00 | 0.03 | 0.090 |
| Stability | Stem density | 2.28 | 0.00 | 0.03 | **0.023** |
| Asynchrony | Species richness | 2.76 | 0.00 | 0.13 | **0.006** |
| Asynchrony | Soil nutrients | -0.84 | -0.19 | -0.04 | 0.401 |
| Asynchrony | $CWM_{SLA}$ | 2.26 | 0.00 | 0.10 | **0.024** |
| Species stability | Species richness | -6.36 | -0.01 | -0.30 | **0.000** |
| Species stability | Soil nutrients | 2.70 | 0.65 | 0.15 | **0.007** |
| Species stability | $CWM_{SLA}$ | -2.72 | 0.00 | -0.12 | **0.007** |
| $CWM_{SLA}$ | Soil nutrients | 7.58 | 106.14 | 0.39 | **0.000** |
| Species richness | Soil nutrients | -11.65 | -63.23 | -0.54 | **0.000** |
| Species richness | Stem density | 5.86 | 0.01 | 0.23 | **0.000** |
| Stem density | Soil nutrients | 3.74 | 663.58 | 0.22 | **0.000** |
| Partial bivariate correlations | | | | | |
| Asynchrony | Species stability | 5.65 | 0.23 | 0.23 | **0.000** |
| Stem density | $CWM_{SLA}$ | 18.57 | 0.61 | 0.61 | **0.000** |

Table S14 The standardized estimates of structural equation models testing the impact of biotic and abiotic factors on community stability for DHS plot. Models are presented in Fig. S5. Significant effects ($P$ < 0.05) are indicated in bold.

| Response | Predictor | Crit. Value | Non-standardized estimate | Standardized estimate | *P*-value |
|---|---|---|---|---|---|
| Stability | Species richness | 0.08 | 0.00 | 0.00 | 0.934 |
| Stability | Asynchrony | 34.71 | 1.29 | 0.75 | **0.000** |
| Stability | Species stability | 19.03 | 0.79 | 0.41 | **0.000** |
| Stability | Soil nutrients | 0.16 | 0.10 | 0.00 | 0.871 |
| Stability | $CWM_{SLA}$ | 1.91 | 0.00 | 0.04 | 0.057 |
| Asynchrony | Species richness | 1.22 | 0.00 | 0.06 | 0.224 |
| Asynchrony | Soil nutrients | -2.14 | -1.69 | -0.11 | **0.033** |
| Species stability | Species richness | -0.29 | 0.00 | -0.01 | 0.770 |
| Species stability | Soil nutrients | 0.98 | 0.72 | 0.05 | 0.327 |
| $CWM_{SLA}$ | Soil nutrients | -0.41 | -42.63 | -0.03 | 0.678 |
| Species richness | Soil nutrients | -0.82 | -11.02 | -0.04 | 0.415 |
| Species richness | Stem density | 17.95 | 0.06 | 0.66 | **0.000** |
| Stem density | Soil nutrients | -0.02 | -3.17 | 0.00 | 0.987 |
| Partial bivariate correlations | | | | | |
| Stem density | $CWM_{SLA}$ | -2.73 | -0.12 | -0.12 | **0.003** |

Table S15 The standardized estimates of structural equation models testing the impact of biotic and abiotic factors on community stability for BN plot. Models are presented in Fig. S5. Significant effects ($P$ < 0.05) are indicated in bold.

| Response | Predictor | Crit. Value | Non-standardized estimate | Standardized estimate | $P$-value |
|---|---|---|---|---|---|
| Stability | Species richness | -1.98 | 0.00 | -0.03 | **0.049** |
| Stability | Asynchrony | 44.76 | 0.69 | 0.59 | **0.000** |
| Stability | Species stability | 52.10 | 0.93 | 0.67 | **0.000** |
| Stability | Soil nutrients | -1.18 | -0.08 | -0.02 | 0.239 |
| Stability | Stem density | 1.15 | 0.00 | 0.02 | 0.251 |
| Asynchrony | Species richness | 0.41 | 0.00 | 0.02 | 0.686 |
| Asynchrony | Soil nutrients | 2.26 | 0.46 | 0.12 | **0.024** |
| Asynchrony | Stem density | 2.41 | 0.00 | 0.15 | **0.016** |
| Species stability | Species richness | -1.13 | 0.00 | -0.06 | 0.260 |
| Species stability | Soil nutrients | 2.23 | 0.41 | 0.13 | **0.027** |
| Species stability | $CWM_{SLA}$ | -2.51 | 0.00 | -0.13 | **0.012** |
| Species stability | Stem density | -1.21 | 0.00 | -0.08 | 0.225 |
| $CWM_{SLA}$ | Soil nutrients | -1.35 | -8.18 | -0.08 | 0.179 |
| Species richness | Soil nutrients | -2.95 | -16.46 | -0.16 | **0.003** |
| Species richness | Stem density | 14.04 | 0.11 | 0.66 | **0.000** |
| Stem density | Soil nutrients | 7.89 | 259.89 | 0.42 | **0.000** |
| Partial bivariate correlations | | | | | |
| Species richness | $CWM_{SLA}$ | 5.73 | 0.25 | 0.25 | **0.000** |
| Asynchrony | Species stability | 3.48 | 0.16 | 0.15 | **0.000** |
